# Diversification of an emerging bacterial plant pathogen; insights from the global spread of *Xanthomonas euvesicatoria* pv. *perforans*

**DOI:** 10.1101/2024.03.22.585974

**Authors:** Sujan Timilsina, Fernanda Iruegas-Bocardo, Mustafa O. Jibrin, Anuj Sharma, Aastha Subedi, Amandeep Kaur, Gerald V. Minsavage, Jose Huguet-Tapia, Jeannie Klein-Gordon, Pragya Adhikari, Tika B. Adhikari, Gabriella Cirvilleri, Laura Belen Tapia de la Barrera, Eduardo Bernal, Tom C. Creswell, Doan Thi Kieu Tien, Teresa A. Coutinho, Daniel S. Egel, Rubén Félix-Gastélum, David M. Francis, Misrak Kebede, Melanie Lewis Ivey, Frank J. Louws, Laixin Luo, Elizabeth T. Maynard, Sally A. Miller, Nguyen Thi Thu Nga, Ebrahim Osdaghi, Alice M. Quezado-Duval, Rebecca Roach, Francesca Rotondo, Gail E. Ruhl, Vou M. Shutt, Petcharat Thummabenjapone, Cheryl Trueman, Pamela D. Roberts, Jeffrey B. Jones, Gary E. Vallad, Erica M. Goss

## Abstract

Emerging and re-emerging plant diseases continue to present multifarious threats to global food security. Considerable recent efforts are therefore being channeled towards understanding the nature of pathogen emergence, their spread and evolution. *Xanthomonas euvesicatoria* pv. *perforans (Xep*), one of the causal agents of bacterial spot of tomato, rapidly emerged and displaced other bacterial spot xanthomonads in tomato production regions around the world. In less than three decades, it has become a dominant xanthomonad pathogen in tomato production systems across the world and presents a model for understanding diversification of recently emerged bacterial plant pathogens. Although *Xep* has been continuously monitored in Florida since its discovery, the global population structure and evolution at the genome-scale is yet to be fully explored. The objectives of this work were to determine genetic diversity globally to ascertain if different tomato production regions contain genetically distinct *Xep* populations, to examine genetic relatedness of strains collected in tomato seed production areas in East Asia and other production regions, and to evaluate variation in type III effectors, which are critical pathogenicity and virulence factors, in relationship to population structure. We used genome data from 270 strains from 13 countries for phylogenetic analysis and characterization of *Xop* effector gene diversity among strains. Our results showed notable genetic diversity in the pathogen. We found genetically similar strains in distant tomato production regions, including seed production regions, and diversification over the past 100 years, which is consistent with intercontinental dissemination of the pathogen in hybrid tomato production chains. Evolution of the *Xep* pangenome, including the acquisition and loss of type III secreted effectors, is apparent within and among phylogenetic lineages. The apparent long-distance movement of the pathogen, together with variants that may not yet be widely distributed, poses risks of emergence of new variants in tomato production.

## Introduction/Main

Emerging and re-emerging plant diseases are a constant threat to global food security [1–3]. Bacterial plant pathogens cause some of the most intractable diseases of crops worldwide [4–7]. Novel emergence and re-emergence of bacterial diseases continue to be reported across the globe and is associated with an upsurge in efforts devoted to understanding the nature of pathogen emergence, spread, and evolution [8–17]. A bacterial plant pathogen that emerged in the last few decades and is of global epidemiological consequences is *Xanthomonas euvesicatoria* pv. *perforans,* one of the causal agents of bacterial spot of tomato [18].

Bacterial spot disease of tomato affects all aboveground plant parts including leaves, stems, flowers and fruit. Under optimal environmental conditions, fruit lesions and/or extensive defoliation can dramatically limit marketable yields and poses a continuous challenge to tomato production [19–21]. Once epidemics are initiated, growers have limited management tools and have relied heavily on copper-based bactericides. However, reliance on copper compounds has led to widespread copper tolerance [22–30].

Alternative bactericides are often costly, provide insufficient control when the weather favors rapid disease development, and rarely improve yields. While historically four taxa have caused this disease, *Xanthomonas euvesicatoria* pv. *perforans* (*Xep*) [31] (syn. *X. perforans* [32, 33]) has emerged rapidly and become a major player on tomato [18, 27, 34–41]. *Xep* was first reported in 1991 in Florida, USA [32] and is now found in all tomato production areas of the world, including regions with no history of the disease [42]. *Xep* has been isolated from tomato seed [32]; therefore, a plausible hypothesis for new outbreaks of *Xep* is pathogen movement with seeds and planting materials [43].

Tomato production is characterized by a high seed replacement rate (99.3%), meaning that growers require seeds each season, which in turn requires large-scale seed production [44]. Tomato hybrid seed production is concentrated in geographic areas where environmental conditions minimize seed contamination by pathogens and seed production costs are low. These seed production regions supply hybrid seeds globally for commercial production of tomato fruits for the fresh market or for processing into tomato products (e.g., sauce, paste, and diced tomatoes). The long-distance movement of seeds poses a high risk for dissemination of seed borne pathogens to commercial tomato production areas. Seedlings are typically grown in transplant facilities and then transplanted into fields for the regional and international transplant markets, potentially amplifying and further disseminating seed-borne pathogens [45].

The success of *Xep* as a pathogen has been attributed to its production of bacteriocins against competing bacterial spot species, rapid genome evolution via recombination affecting the chromosome, and introduction of genes via horizontal gene transfer that contribute to fitness in tomato fields [46–52]. Distinct genetic lineages of *Xep*, each with unique patterns of allelic variation among core genes (genes present in all strains), were identified in fresh market and processing tomato production fields in the United States [47, 50, 51, 53, 54]. Additional lineages of *Xep* were found in Nigeria, Iran, Italy, and the Southwest Indian Ocean islands [42, 48, 55].

*Xanthomonas perforans* strains, like other xanthomonads, acquire nutrients through colonization of compatible hosts. The type III secretion system (T3SS) and type III effector (T3E) proteins are critical for suppression of host defenses and virulence by *Xep* [56]. Effector content varies among *Xanthomonas* species and distinct lineages of *Xep* have distinguishable effector content [23, 48, 50, 57, 58]. Strains of *Xep* isolated in the 1990s were limited to tomato [59], but now strains of *Xep* are causing bacterial spot disease of pepper [50, 58, 60]. Host range expansion was attributed, in part, to loss of effectors that act as avirulence factors in pepper and other genomic changes as a result of recombination with other *Xanthomonas* lineages, including pepper pathogenic *X. euvesicatoria* pv. *euvesicatoria*. Effector variation may cause differences in disease epidemiology in addition to host range [49, 57, 61]. For example, wildtype strains with the acquired effector XopJ2 showed three times faster spread in the field than isogenic mutant strains without the effector [51].

Emerging pathogens may show limited genetic variation if they experienced a bottleneck during the ecological and evolutionary processes that often precede emergence (e.g., host jump or introduction event) [62]. *Xep* appears genetically diverse but it is not known how this variation is structured across global tomato production regions. The first objective of this work was to determine if different tomato production regions contain genetically distinct *Xep* populations. Second, we asked if there was evidence for long-distance pathogen dissemination, as would be indicated by genotypes shared among distant regions. Specifically, we obtained strains from tomato seed production areas in East Asia and determined if they resembled strains from fruit production fields elsewhere in the world, which would be expected if strains are being disseminated in seeds. Third, we estimated the timing of *Xep* population expansion relative to its first report in 1991. Finally, we evaluated T3SE content and allelic variation in the context of geography and core genome variation as a proxy for genetic variation in virulence. Overall, we found extensive genetic diversity within *Xep;* genetically similar strains in distant geographic regions, inclusive of seed production regions; evidence of diversification prior and subsequent to the first report of emergence; and lineage-specific T3SE repertoires. Together, these results illustrate the capacity for this pathogen to rapidly evolve and strongly support the potential for intra- and intercontinental movement of pathogens in tomato production systems.

## Results

### *X. euvesicatoria* pv. *perforans* strains from seed and commercial fruit production areas

A total of 270 *Xep* genomes from 13 different countries – representing seed and fruit production – were used in this study (Table 1). We generated new genome sequence data for 153 strains (S1 Table; NCBI BioProject PRJNA941448). *Xep* strains were differentiated from other tomato-pathogenic xanthomonads using a real-time qPCR assay that specifically amplifies the *hrcN* (*hrpB7*) gene in *Xep* [63] and inoculated on tomato cv. ‘Bonny Best’ to confirm pathogenicity. Strains from China, Thailand, and Vietnam were collected from seed production areas (n = 31) and all other strains (n = 239) were collected in commercial fruit production areas from Australia, Brazil, Canada, Ethiopia, Iran, Italy, Mexico, Nigeria, South Africa, and the United States. Within the US, strains were collected from seven different states in the Midwest and Southeast, including strains collected since 1991 from Florida.

**Table 1.** *Xanthomonas euvesicatoria* pv. *perforans* strains used in this study.

| Country | Locality | Year | Strain (original name, if applicable) |
| --- | --- | --- | --- |
| Australia [57] | Queensland | 2015 | Aus3, Aus7, Aus14 |
|  |  | 2016 | Aus5, Aus10, Aus11 |
|  |  | 2017 | Aus1, Aus15, Aus16 |
| Brazil [39] | São Paulo | 2011 | Bzl1 (2011-107), Bzl2 (2011-132) |
|  | Goiás | 2012 | Bzl3 (2012-08) |
|  | Goiás, São Paulo | 2013 | Bzl5 (2013-16), Bzl6 (2013-42) |
|  | Goiás, Minas Gerais | 2014 | Bzl7 (2014-10), Bzl8 (2014-17) |
|  | Minas Gerais | 2015 | Bzl10 (2015-53), Bzl11 (2015-56) |
|  | Goiás | 2016 | Bzl13 (2016-08) |
|  | Goiás | 2017 | Bzl14 (2017-21) |
| Canada | Ontario | 2016 | 4A, 4D, 12A, 14A |
| China |  | 2016 | CHI-3, CHI-5, CHI-6, CHI-7, CHI-8, CHI-10, CHI-12, CHI-15, CHI-18 |
| Ethiopia [36] |  | 2011 | ETH5, ETH11, ETH21, ETH25, ETH33 |
| Iran [41] |  | 2013 | K41, F210, F215, TOM801, TOM816 |
| Italy [64] |  | 2011 | 1P6S1, 2P4S1, 2P4S1D, 2P6S1, 1P4S1D |
| Mexico |  |  | Mexico-1, Mexico-3, Mexico-LT1, Mexico-LT3, Mexico-LT5 |
| Nigeria [37, 65] |  | 2014 | NI-1, NI-2, NI-4, NI-7, NI-12, NI-13 |
|  |  | 2015 | KS3, KS5, KS9, KS28 |
| South Africa | Pretoria |  | X2-B14, X10-B85, X59-BD1351, X47-BD167 |
| Vietnam |  |  | SEA-3, SEA-5, SEA-21, SEA-23 |
| Thailand |  | 2016 | THA-8, THA-14, THA-40, THA-45, THA-54, THA-72,<br>THA-81A, THA-100, THA-112, THA-116, THA-119,<br>THA-120, THA-126, THA-127, THA-128, THA-132,<br>THA-135, THA-157A |
| United States | Alabama [66] | 1996 | Xp1861 |
|  | Indiana [25] | 2016 | 16-1165A1, 16-1181-2, 16-1182A, 16-1184A, 16-1187A,<br>16-1205A, 16-1402A, 16-974C, 16-990A, 16-990C |
|  |  | 2014 | 14-463-1A |
|  | Florida [32,<br>33, 43, 47, 58,<br>67] | 1991 | XV0938, Xp91-118, Xp894, Xp909, Xp1183 |
|  |  | 1992 | Xp1118, Xp1144 |
|  |  | 1993 | Xp1241, Xp1268, Xp1275 |
|  |  | 1994 | Xp1550, Xp1564 |
|  |  | 1995 | Xp1797, Xp1805 |
|  |  | 1996 | Xp1856 |
|  |  | 1997 | Xp1912 |
|  |  | 1998 | Scott-1, Xp1920 |
|  |  | 2006 | Xp1-5, Xp1-6, Xp3-12, Xp3-15, Xp3-16, Xp3-8, Xp4-20,<br>Xp5-14, Xp5-6, Xp5-9, Xp7-12, Xp8-16, Xp9-5, Xp10-13, Xp11-2, Xp15-11, Xp17-12, Xp18-15 |
|  |  | 2007 | Xp4B |
|  |  | 2010 | Xp2010 |
|  |  | 2011 | GEV485 |
|  |  | 2012 | GEV839, GEV872, GEV893, GEV904, GEV909,<br>GEV915, GEV917, GEV936, GEV940, GEV968,<br>GEV993, GEV1001, GEV1026, GEV1044, GEV1054,<br>GEV1063 |
|  |  | 2013 | TB6, TB9, TB15 |
|  |  | 2015 | GEV2047, GEV2048, GEV2049, GEV2050, GEV2052,<br>GEV2055, GEV2058, GEV2059, GEV2060, GEV2063,<br>GEV1989, GEV1991, GEV1992, GEV1993, GEV2004,<br>GEV2009, GEV2010, GEV2011, GEV2013, GEV2015,<br>GEV1911, GEV1912, GEV1913, GEV1914, GEV1915,<br>GEV1916, GEV1917, GEV1918, GEV1919, GEV1920,<br>GEV1921 |
|  |  | 2016 | GEV2065, GEV2067, GEV2072, GEV2087, GEV2088,<br>GEV2089, GEV2097, GEV2098, GEV2099, GEV2108,<br>GEV2109, GEV2110, GEV2111, GEV2112, GEV2113,<br>GEV2114, GEV2115, GEV2116, GEV2117, GEV2118,<br>GEV2119, GEV2120, GEV2121, GEV2122, GEV2123,<br>GEV2124, GEV2125, GEV2126, GEV2127, GEV2128,<br>GEV2129, GEV2130, GEV2132, GEV2133, GEV2134,<br>GEV2135 |
|  | Louisiana [24] | 2013 | mli-2 |
|  | North<br>Carolina [27] | 2015 | NC-14, NC-47, NC-67, NC-101, NC-112, NC-204 |
|  |  | 2016 | NC-242, NC-252, NC-282, NC-289, NC-350, NC-373,<br>MRS-30P-011 |
|  | South<br>Carolina [43] | 2016 | GEV2407, GEV2408, GEV2384, GEV2388, GEV2389,<br>GEV2390, GEV2391, GEV2392, GEV2393, GEV2396,<br>GEV2397, GEV2399, GEV2400, GEV2403, GEV2410,<br>GEV2420 |
|  | Ohio [53] | 2017 | SM-1806, SM-1807, SM-1808, SM-1809, SM-1810, SM-1811, SM-1812, SM-1813, SM-1814, SM-1815, SM-1828, SM-1829, SM-1830, SM-1831 |

### Genomic diversity in *X. euvesicatoria* pv. *perforans*

To examine genetic diversity in the core genome, we curated a set of 887 genes that were present in all 270 *Xep* genomes based on IMG/JGI gene annotation. The aligned sequence length of concatenated core genes was 617,855 bp, which contained 14,427 polymorphic sites after removing ambiguous nucleotides and any alignment gaps (S1 Data). Maximum likelihood phylogenetic analysis showed a distinct and especially diverged lineage of 11 strains from Nigeria and Thailand (S1 Figure), that included a previously defined atypical strain – NI1 – from Nigeria [48]. Grouping strains by state within the United States and country elsewhere produced an FST [68] of 0.66. The number of core gene SNPs by geographic location represented by more than one strain ranged from 15 to 5929 (S2 Table). Nucleotide diversity (average number of differences among sequences) ranged from 3 to 1287, with both extremes in nucleotide diversity coming from the Midwestern U.S., Ohio and Indiana respectively (S2 Table). Tajima’s D [69] by geographic location ranged from –2.0 to 1.7 (S2 Table).

Phylogenetic analysis of core SNPs, followed by correction of branch lengths for recombination, showed diversifying lineages of *Xep* (Figure 1; S1 Figure). After excluding a particularly diverged lineage of 11 strains (S1 Figure), ClonalFrameML [70] estimated an overall ratio of recombination rate to mutation rate (R/theta) of 0.60, with recombination causing approximately seven times more base changes than mutation (delta = 231; nu = 0.05). There were an estimated 221 recombination events that affected more than 96 Kbp in terminal branches and 494 recombination events detected in internal branches encompassing 190 Kbp.

**Figure 1.**
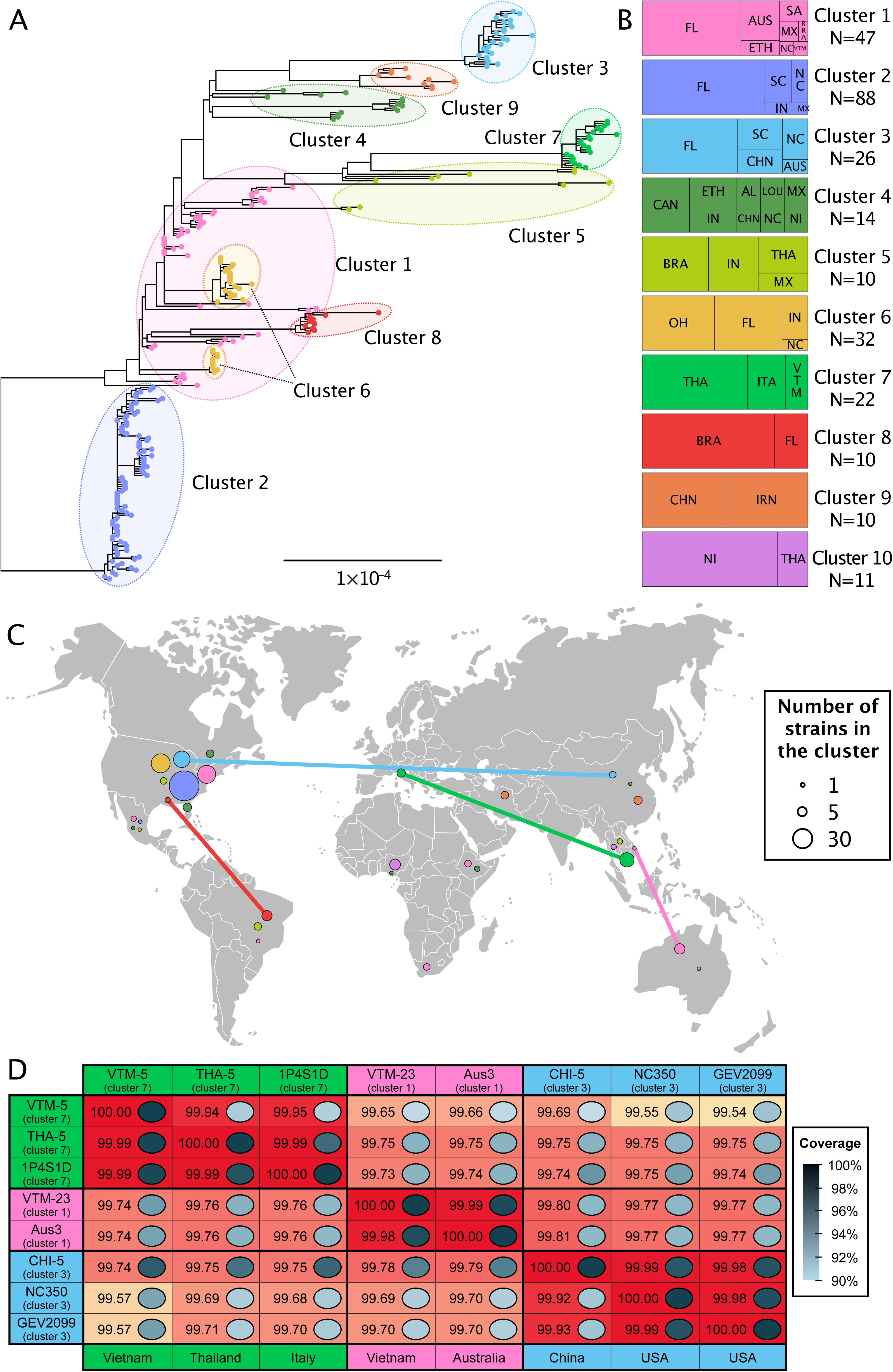
Population structure of *Xanthomonas euvesicatoria* pv. *perforans* strains collected from tomato production regions. (A) Maximum likelihood phylogenetic tree of 259 *X. euvesicatoria* pv. *perforans* strains constructed with nucleotide sequences from 887 core genes, corrected for recombination by ClonalFrameML. Tips are colored according to clusters identified by hierBAPS. Cluster 10 strains (n=11) were genetically distant and excluded from the tree (see S1 Figure). Nucleotide alignment is available as S1 Data. (B) Distribution of 270 strains in each hierBAPS cluster by country or state of collection. Abbreviations are as follows: AUS – Australia; BRA – Brazil; CAN – Canada; CHN – China; ETH – Ethiopia; FL – Florida, USA; IN – Indiana, USA; IRN – Iran; ITA – Italy; LOU – Louisiana, USA; MX – Mexico; NC – North Carolina, USA; NI – Nigeria; AL – Alabama; OH – Ohio, USA; SA – South Africa; SC – South Carolina, USA; THA – Thailand; VTM – Vietnam. (C) Map showing distribution of clusters by country of collection. Countries connected by lines show selected instances of genetically similar strains collected in different countries. (D) Pairwise comparison of whole genome average nucleotide identity (ANIb) of selected strains shows high identity between strains isolated from different continents. For each comparison, genome coverage is shown by grayscale in boxes, scale shown to the right. Values for each comparison are for genomes in rows when compared to genomes in columns. See S3 Table for additional ANI output.

To summarize population structure based on core gene SNPs, we used hierBAPS [71], which assigned individual strains to 9 clusters using allele frequencies (Figure 1; S1 Table). This analysis excluded the 11 highly diverged strains from Nigeria and Thailand, which we designated as cluster 10. F_ST_ among clusters was 0.80. In some cases, clusters corresponded to phylogenetic lineages, including clusters 2, 3, 7, 8, and 9 (Figure 1). The remaining clusters were polyphyletic, encompassing multiple diverged clades or individual strains. Nucleotide diversity within clusters ranged from 13.8 to 675.9 and Tajima’s D from –2.6 to 1.0. Presence-absence gene variation in the pangenome largely paralleled the phylogenetic diversity of core genes in that polyphyletic clusters 1, 4, and 5 also showed the most variation in gene content (S2 Figure).

### Geographic distribution of *X. euvesicatoria* pv. *perforans* core gene clusters

Cluster 1 encompasses genetically diverse strains from seven countries, including most of the strains from Australia, all four strains from South Africa, and one strain from Southeast Asia (Figure 1B). All USA strains assigned to cluster 1 were isolated in or before 2006 from Florida except for one strain from North Carolina. Cluster 2 contains 88 strains from the United States and one from Mexico, while Cluster 3 includes strains isolated from Florida, North and South Carolina, China, and Australia. Cluster 4 encompasses multiple lineages of strains from the United States, Canada, Ethiopia, China, and Nigeria. Cluster 5 is polyphyletic with diverged strains from three continents. Cluster 6 was isolated only within the United States from Florida, Indiana, North Carolina, and Ohio. Cluster 7 is a monophyletic group of strains from Southeast Asia and Italy. Cluster 8 is another monophyletic group found only in Brazil and Florida. Cluster 9 includes two clades of strains, one from China and the other from Iran and Nigeria. Cluster 10 comprises the atypical strains from Nigeria and similar strains from Thailand. Most countries contained strains from more than one core gene cluster (Figure 1C).

Clusters 1, 3, 4, 5, 7, 9, and 10 contain strains isolated from both seed production and commercial fruit production regions, whereas strains in clusters 2, 6, and 8 were only isolated from commercial fruit production regions. Some strains found on different continents were nearly identical in core gene sequences (Figure 1D) with very high average nucleotide identity. Strains in cluster 1 from Australia differed by 6 to 10 SNPs in more than 617 Kbp of core gene sequence from strain VTM-23 from Vietnam. Pairwise average nucleotide identity (ANIb) between VTM-23 to Aus3 was 99.99% with alignment fraction of 0.998 (Figure 1D). Strains from the USA had up to 99.87 ANIb with strains from Australia and Vietnam (S3 Table). A different strain from Vietnam, VTM-5 in cluster 7, had as few as four SNPs in the core genome when compared to strains from Italy and ANIb of 99.95% to Italian strain 1P4S1D (Figure 1D). Likewise, strains collected in a seed production region in China had ANIb up to 99.99% with strains from Florida and North Carolina. We also found similar strains between Brazil and USA, for example Bzl-10 (Minas Gerais) and Xp3-8 (Florida) had greater than 99.9% ANI (S3 Table). Other strains were similar between countries in core genes only after correction for recombination.

### Timing of *X. euvesicatoria* pv. *perforans* lineage emergence

We used the years of strain collection to estimate the timing of diversification of our sample of *Xep*, excluding the cluster 10 strains. We inferred dated phylogenies using whole genome alignments with inferred recombinant sites removed by Gubbins [72]. Due to recombination with other *X. euvesicatoria* lineages, we did not include an outgroup (S1 Figure, part B; [48]). Sampling year was significantly correlated with root-to-tip distance (R^2^ = 0.20 for the whole genome alignment, *P* < 1×10^-4^, S3 Figure). The root inferred by the BactDating R package [73] was placed between strains isolated in Florida in 1991. The most recent common ancestor (MRCA) of all strains was dated to 1884 (95% HPD: 1655– 1966). Notably, strains that were isolated in the early 1990s, when *Xep* was first detected in U.S. tomato production [32, 37], represented multiple lineages (Figure 2). The MRCA of the clade representing core gene clusters 1, 2, 6, and 8 (including strains from USA, Brazil, and Mexico) was dated to 1980 (95% HPD: 1967–1987). A major clade, encompassing strains in clusters 3, 4, 5, 7, 9, which were collected in Africa, the Americas, Asia, Australia, and Europe, did not have a significant temporal signal across the clade. We repeated the analysis with BEAST, which inferred a different rooting. The tree inferred by BEAST placed the root between two strains isolated in 2011 from Brazil and all other strains (S4 Figure, part B). The MCRA of the BEAST tree was dated to 1868 (95% HPD: 1862–1919), which was similar to the root date estimated using BactDating (S4 Figure).

**Figure 2.**
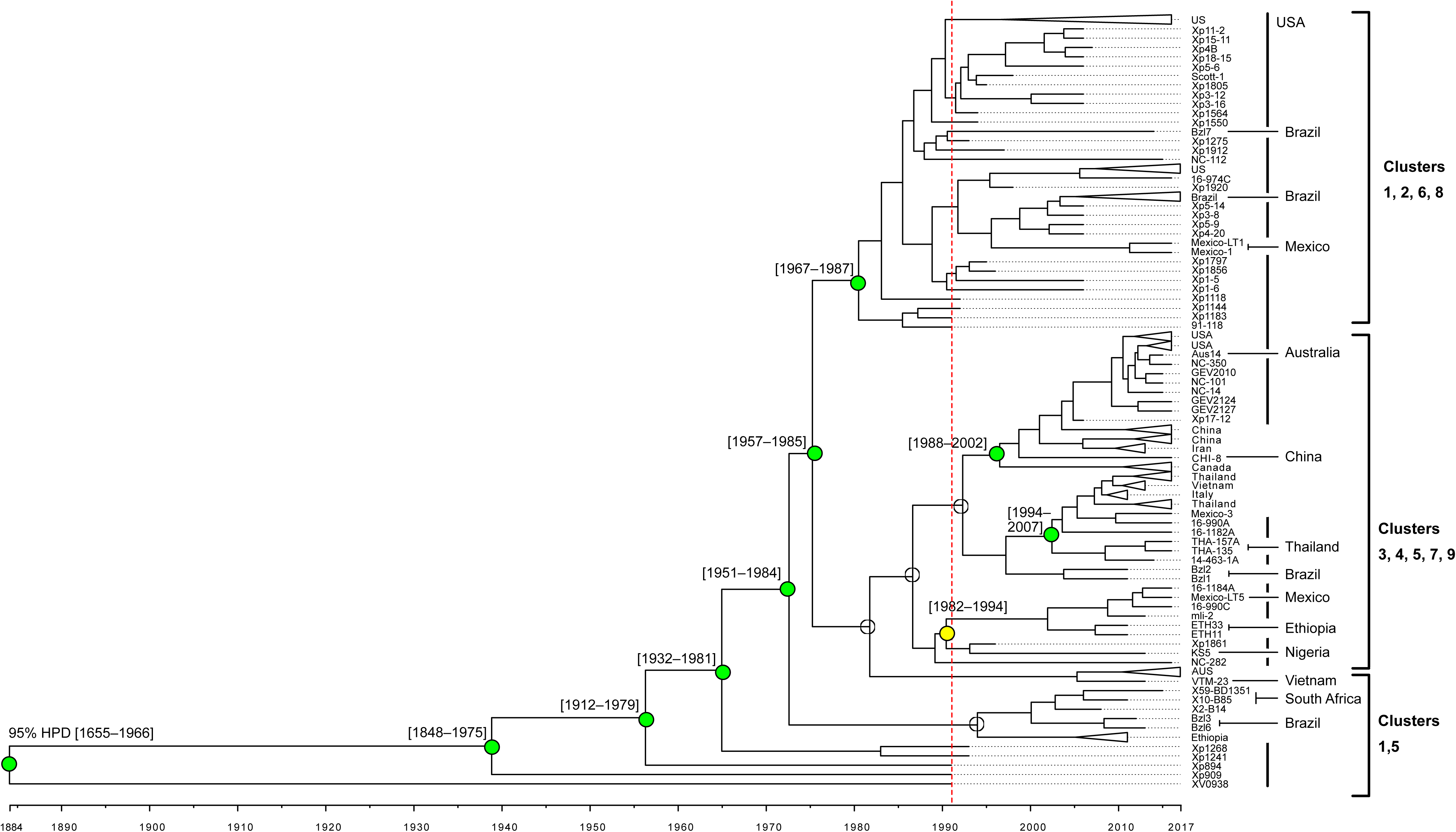
Dated phylogeny of 259 *X. euvesicatoria* pv. *perforans* strains. BactDating analysis estimated an approximately 130-year history for *Xep* strains in core gene clusters 1 through 9 (Figure 1). Red dotted line indicates the first documented isolations in 1991. Internal nodes were collapsed for clades containing strains from a single country with branch tips indicating country or strain (for full tree see S4 Figure). Vertical line to the right of tip labels indicates strains from USA; other countries are labeled. Temporal signal was assessed using Phylostems and results are shown for major nodes (for full results see S3 Figure). Empty circles indicate no significant temporal signal. Colored circles indicate nodes with statistically significant temporal signal based on adjusted R^2^ values: green – 0.13-0.19; yellow – 0.45. The 95% highest posterior density (95% HPD) of date estimates for major nodes with significant temporal signals are shown in brackets.

### Type III effector content

We detected 32 predicted type III effectors in our collection of 270 strains (S5 Figure, S4 Table). The diversity in amino acid sequences of predicted effectors ranged widely from a single conserved allele to 8 or more alleles per locus (Figure 3). None of the effectors were present and intact in 100% of our genomes, in part due to our analysis of draft genomes. The following effector genes were present in more than 95% of strains and can be considered “core effectors”: *avrBs2*, *xopF1*, *xopF2*, *xopI*, *xopM*, *xopQ*, *xopS*, *xopV*, *xopX*, *xopAE*, *xopAK*, *xopAP*, *xopAU*, and *xopAW*. The genes for *xopD*, *xopE1*, and *xopN* were present in some form in all genomes but more than 5% of strains contained a contig break within the gene. A closer examination of *xopD* by PCR and Sanger sequencing showed this to be an assembly issue due to the repeats within the gene. Effectors at low frequency in our *Xep* strains (<25%) were *xopE3*, *xopAD*, *xopAJ*, *xopAO*, and *xopAQ*. Transcription activator-like (TAL) effectors typically do not assemble in draft genomes due to their characteristic repeat sequences, but there were BLAST hits to previously described TAL effectors in 65 strains. We Sanger sequenced the TAL effector gene in strain 2P6S1 collected in Italy (NCBI accession number OQ588696), which confirmed that it had the same repeat variable diresidues as PthXp1 reported in *Xep* strains from Alabama [50]. Previous phenotyping and sequencing indicated that the strain isolated in Louisiana, USA origin has AvrHah1 [16].

**Figure 3.**
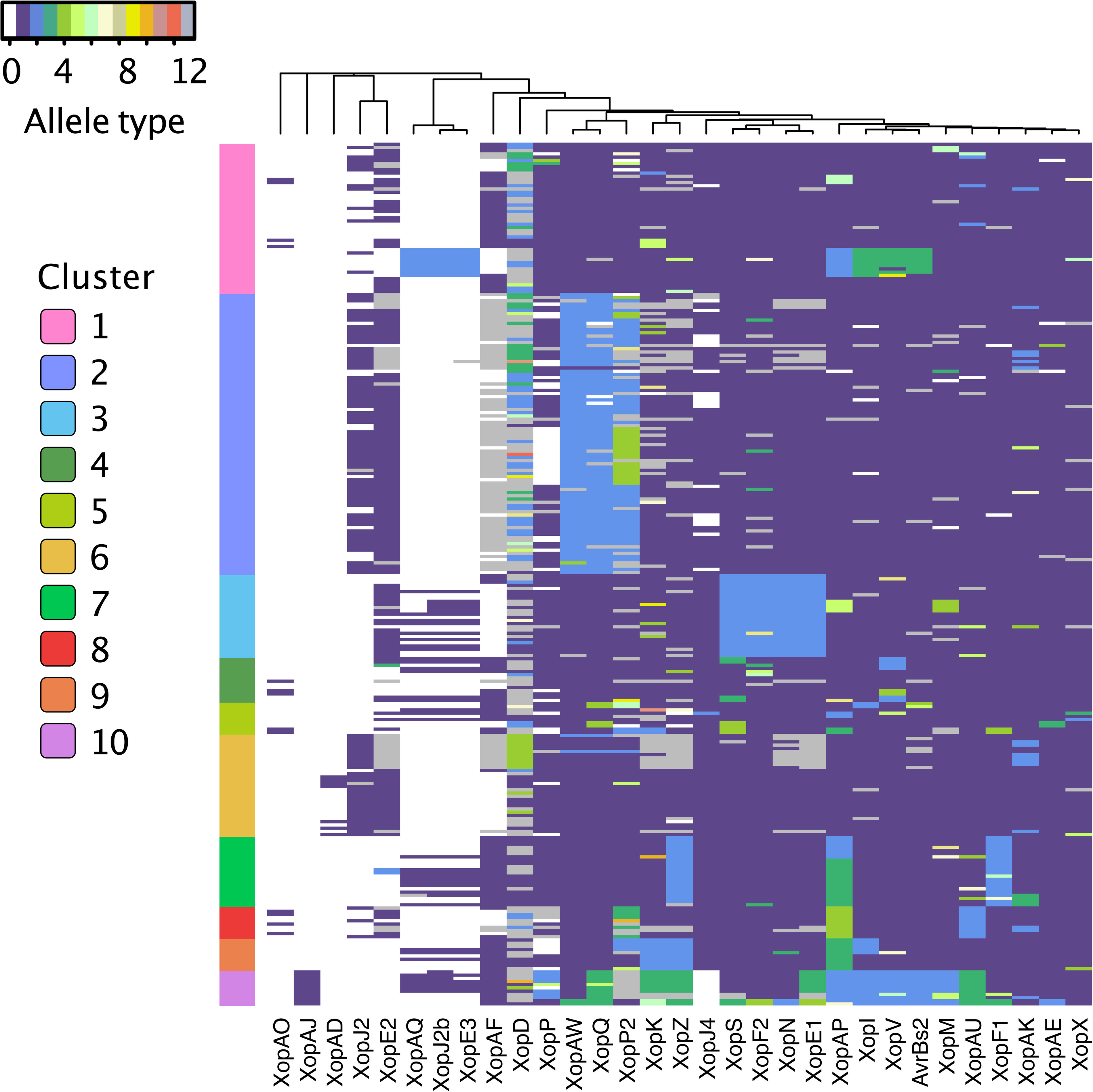
Variation in type III effectors (Xop proteins) in *Xanthomonas perforans*. Type III effectors are in columns and 270 *X. euvesicatoria* pv. *perforans* strain in rows. Effector status is shown by allele type: absence is indicated by allele type 0 (white), while the most frequent allele observed when the effector is present is allele type 1 (purple), second most frequent is allele type 2 (blue), and so on. Putative pseudogenized effectors are shown as allele 13 (gray). The order of columns was determined by hierarchical clustering analysis, placing similarly distributed effectors adjacent to each other. Genomes showing BLAST hits to TAL effector(s) are indicated in Supporting Information Table S3 and not shown in heatmap. *X. euvesicatoria* pv. *perforans* strains (rows) are organized by core gene cluster.

The T3E effector XopAF (AvrXv3), which is targeted by the tomato resistance gene *Xv3* [74], was missing or pseudogenized in 64% of strains. Most strains examined from the United States did not have a complete copy of this gene, whereas it was intact in many strains collected in Asia and Africa. The gene for XopJ4 (AvrXv4), recognized by resistance gene *RXopJ4* from *S. pennellii* [75], was present in 88% of strains and absent in all cluster 10 strains and 19 of 88 cluster 2 strains. XopJ2 (AvrBsT), which elicits an HR in pepper but increases virulence in tomato [49], was present in less than half of strains examined (43%) and overwhelmingly in strains from the United States. A homolog of XopJ2, recently designated XopJ2b [76], was present in 50 strains, including two strains from Australia that carried both copies of XopJ2 (S4 Table).

We tested for evidence of positive selection in T3E by estimating synonymous and non-synonymous (dN/dS) substitution rates using a Bayesian approach for detecting pervasive selection (FUBAR, [77]) and maximum likelihood approach for detecting episodic selection (MEME, [78]). We found evidence of pervasive positive selection affecting at least one amino acid in AvrBs2, XopD, XopE1, XopF2, XopK, XopM, XopP and its paralog XopP2, XopQ, XopS, and XopAQ (S4 Table). We found evidence of episodic selection affecting at least one amino acid in XopF2, XopK, XopP, XopP2, XopQ, XopV, and XopAP (S4 Table).

We defined the effector profile of each strain as the predicted presence or absence of each effector and its allelic state, excluding TAL effector hits. Grouping effector profiles according to core gene cluster revealed that allelic variation of effectors often paralleled core genome variation (Figure 3). For example, specific alleles of effectors XopAW, XopQ, and XopP2 were mostly limited to strains in cluster 2. Cluster 3 strains carried unique alleles for effectors XopF2, XopS, XopN, and XopE1, and strains from cluster 7 shared unique alleles for effectors XopF1 and XopZ. Strains from highly diverged cluster 10 had rare alleles in many effectors, and it was the only cluster in which effector XopAJ was found (Figure 3). To visualize variation among strain effector profiles independent of core gene clusters, we transformed dissimilarities between profiles into distances represented in a two-dimensional plot and defined eight effector profile clusters (S6 Figure, part A). A lack of low frequency effectors characterized effector cluster A, containing 188 strains from 11 of 13 countries (S6 Figure). The remaining effector clusters were defined by the presence of one to three low frequency effectors (S7 Figure). While most effectors were found in multiple countries and continents (S6 Figure), populations in Brazil, Ethiopia, Nigeria, Thailand, South Africa, and the United States contained low frequency effectors that were not widely distributed.

### Copper resistance genes

Xanthomonads, including *Xep*, have acquired genes conferring copper tolerance, likely in response to exposure to copper-based bactericides [30, 79–81]. In *Xep*, copper tolerance is conferred by an operon containing the copper resistance genes *copA* and *copB*, and regulator *copL* (*copLAB*) [80]. BLAST analysis showed that these genes were present in 73% of the genomes in our sample (S5 Table). Copper resistance genes are prevalent in the USA; only the genomes from strains isolated from Florida in the early 1990s and a strain from Louisiana lacked *copLAB*. The genes were also missing in the genomes of a few strains from Australia (1), Brazil (2), Ethiopia (2), Mexico (1), and Vietnam (2). In contrast, the genes were absent in all genomes of all strains from Nigeria, China, Iran, Italy, and Thailand.

## Discussion

Emerging plant pathogens have the potential for global outbreaks, exacerbated by complex trade networks. Hybrid tomato production relies on international breeding and production chains with a global network to deliver seeds to growers. Global trade associated with vegetable seed production provides a pathway for global spread of pathogens, with quantities traded that challenge even strong phytosanitary measures [82, 83]. Over 100 countries import seeds of tomatoes and other vegetables; for example, 11.7 million kg of vegetable seed were imported to the USA in 2019, with China being the biggest supplier at 2.4 million kg [84]. *Xanthomonas* species can infest pepper and tomato seed [85], and *Xep* has been isolated from tomato seed [32, 86], supporting the hypothesis that seeds can be a source of inoculum for bacterial spot outbreaks [87, 88]. Thirty years after its first report, *Xep* has been identified in tomato production areas around the world [18]. Our results showed extensive genetic diversity in the pathogen, but also genetically similar strains in distant tomato production regions. Furthermore, we found genetically similar strains in seed production and fruit production regions on different continents, as would be expected if the pathogen was being moved in shared production chains. Dated phylogenies indicate multiple waves of diversification of the *Xep* population, before and since its first detection in 1991. Variation in gene content confirms that *Xep* acquired and lost type III effectors during its diversification, which will continue to challenge sustainable management of tomato bacterial leaf spot [49, 66].

Using our broad strain collection, we found *Xep* variants in seed production regions in Asia that were previously reported in Australia, Italy, Nigeria, and the United States [43, 47, 48, 57]. Strains from Italy were nearly identical in core genes and very similar in accessory genomes to strains collected from Thailand and Vietnam, both major seed production regions. The atypical bacterial spot strains from Nigeria, recently designated as race T5 [21], were genetically similar to strains from Thailand. A recently described variant of *Xep* in Florida [cluster 3; [43, 47]], which was also found in Australia [57], was similar in the core genome to strains found in China; however, these strains showed divergence in the pangenome, consistent with accessory genome evolution in emergent populations. Beyond previously described variants, we found strains in Iran that were closely related to strains from China; multiple instances of genetic similarity between strains from North America and Ethiopia; and highly similar strains shared between USA and Brazil, USA and Australia, and between Australia and Vietnam. Given the variation of *Xep* across our sample, genetic similarity in core genes and gene content across continents is strong evidence of international dissemination. Genetically similar strains of bacterial spot pathogens *X. euvesicatoria* pv. *euvesicatoria* and *X. hortorum* pv. *gardneri* collected from different continents similarly suggest intercontinental dissemination in tomato and pepper seed [15, 58, 89]. Whole genome analysis of *X. hortorum* pv. *pelargonii* strains from a 2022 epidemic of bacterial blight of geranium in the USA showed zero to seven chromosomal SNPs among isolates of the emergent strain that was distributed to multiple states in plant cuttings [9, 90].

Other *Xep* genotypes indicated a more limited distribution. We did not find core gene cluster 2 strains in the seed production regions sampled (China, Thailand, Vietnam), while this lineage was highly represented in our USA sample. To date, strains in this cluster have been found only in the southeastern and midwestern USA [43, 47, 50, 53, 54] and Mexico. Seedling nurseries in the southeast USA produce tomato transplants for growers in multiple states. Interstate movement of strains on seedlings is likely responsible, at least in part, for disseminating genetically similar strains to different states [43, 53]. We previously reported extensive recombination with *X. euvesicatoria* pv. *euvesicatoria* in cluster 2 strains [47] and this cluster had a diverse accessory genome, perhaps suggesting that it has a larger geographic distribution than represented in our sample, which is biased towards the USA.

The *Xep* strains we examined from the USA were assigned to core gene clusters 1, 2, 3, 4, 5, 6, and 8, representing several distinct genetic lineages. To better understand the initial emergence of *Xep*, we used calibrated phylogenies to examine the timing of lineage divergence. International trade in F1 hybrid tomato seed surged in the second half of the 20^th^ century, after the first hybrid tomato cultivars were released by 1940 [91, 92]. There was a 300-fold increase in hybrid tomato seeds exported from Asia between 1962 and 1977 [93, 94] and subsequent rapid growth in tomato production. Our analyses estimate the most recent common ancestor of our sample to ∼150 years ago, while the major ancestral lineages diverged during or after the early expansion in the hybrid seed trade. We hypothesize that the emergence and geographic distribution of lineages may be associated with the multinational structure of tomato breeding and seed production, in which parental lines and geographic locations of seed production change over time [95].

The roles of T3E in pathogenicity and virulence make them important members of both the core and accessory genomes. We found up to 16 putative core effector genes, most of which exhibited allelic variation. The impact of allelic variation in *Xep* effectors on pathogen fitness, if any, is unknown. Low frequency effectors were found across core gene clusters, suggesting acquisition of new effectors and their exchange among *Xep* lineages. For example, some strains in clusters 3, 4, and 5 from the United States, Canada, and Mexico carried the same alleles of low frequency effectors XopAQ and XopE3 as strains from Asia, Nigeria, and Italy. BLAST analysis suggested the geographically widespread presence of transcription activation-like (TAL) effectors in *Xep*. Both TAL effectors described in *Xep*, *avrHah1* and *pthXp1*, are associated with increased disease severity on tomato [16, 50]. Acquisition of T3Es could increase the fitness of *Xep* relative to other bacterial spot pathogens and cause more damaging disease outbreaks [49, 51, 66].

The release of new plant varieties that carry disease resistance genes can have dramatic effects on pathogen population structure due to selection to overcome host resistance [96–98], and we have previously reported on the loss of function of effector AvrXv3 (XopAF) across lineages [27, 53, 66]. Examination of T3E content at a global scale puts variation previously observed in Florida into a larger context. XopAF was present in strains collected in the 1990s (cluster 1), but absent or non-functional in most strains from Florida, Indiana Ohio, and North Carolina, USA [23, 25, 27, 53, 66]. Here, we found that *xopAF* was intact in many strains from seed production areas, suggesting strong selection for loss of function in commercial fruit production. The effector XopJ4 is a possible resistance target [66], but Klein-Gordon et al. [23] reported that it was missing from 3.2% of Florida strains collected in 2017 and, here, we found that it was absent in one North Carolina and 20 Florida, USA strains. All strains collected outside the USA contained *xopJ4*, except for cluster 10 strains. Another XopJ family member, *xopJ2*, is a virulence factor in tomato [49, 51]. It is common in North America, particularly in cluster 2 and 6 strains, but absent or infrequently detected in *Xep* populations elsewhere. An alternative form of this effector, recently described as XopJ2b [76], is more common in strains from outside North America.

Bacterial spot is a destructive disease in areas where tomatoes are grown under humid conditions and growers in the USA have relied heavily on copper bactericides to manage this disease. In response, *Xep* strains have developed copper tolerance [99]. Most strains isolated from Florida in the 1990s lacked the *copLAB* genes, but they are now common in strains collected in the USA. A recent study of Florida strains found that these copper resistance genes are more frequently present on the chromosome than on a plasmid, suggesting selection for vertical inheritance of copper tolerance [30]. In contrast, strains from other countries lacked copper resistance genes, indicating little or no local selection for the acquisition of *cop* genes.

In summary, we found strong evidence for intercontinental movement of *Xep*, consistent with the international nature of tomato breeding and hybrid tomato seed production. We also found notable diversity in our global sample of *Xep*, including in seed production regions, and multiple variants of *Xep* that do not appear to be widely distributed. The genomic diversity of *Xep* in seed and fruit production regions creates the opportunity for recombination among strains and subsequent dissemination of high fitness variants of *Xep*.

## Materials and Methods

### Bacterial strains, genome sequencing, and assembly

*Xep* strains were collected from 13 different countries (Table 1; S1 Table). Strains from the United States were collected from seven states between 1991 to 2016 and comprised 181 strains. The remaining 89 strains were collected from Canada, Mexico, Brazil, Italy, Ethiopia, Nigeria, South Africa, Iran, China, Thailand, Australia, and Vietnam. Strains from China, Thailand, and Vietnam were collected from fields designated for production of tomato seed for the global market. Strains from Brazil were obtained from both staked fresh-market and processing tomato commercial fields. Strains from Italy were isolated from tomato pith in greenhouse tomato showing wilting symptoms [64, 100]. Strains from South Africa were collected from commercial seed lots. Strains from Nigeria were obtained from fields cultivated for both subsistence and commercial purposes. Strains from other countries were collected from fields designated for commercial fruit production.

A total of 270 *Xep* genome sequences were used during this study (S1 Table). Draft and whole genomes of 117 strains were generated and published previously [43, 47, 48, 57, 58, 67, 100]. The remaining 153 strains were sequenced for this study using Illumina platforms. Genomic DNA was extracted from single colony cultures grown for 24-hr in nutrient broth using the Wizard Genomic DNA Purification Kit (Promega, Chicago, IL) following manufacturer instructions. Genomic libraries for sequencing were prepared using the Nextera DNA library preparation kit from Illumina (Illumina, San Diego, CA). Sequencing was performed at the Interdisciplinary Center for Biotechnology Research, University of Florida, using an Illumina MiSeq to generate 250 bp paired end reads for each strain. Additional genomic sequence data were generated for five strains for the ANI analysis (S1 Table, part B). Genomic DNA was extracted using the above methods except that extracted genomic DNA was sent to SeqCenter (Pittsburg, PA) for sequencing with Illumina NovaSeq 6000, producing 150 bp paired end reads.

Raw reads were trimmed of adapters and paired with Trim Galore (https://github.com/FelixKrueger/TrimGalore)[101], then assembled into contigs with Spades version 3.10.1 [102], with k-mers 21, 33, 55, 77, 99, and 127 with read error correction and “--careful” switch. Reads were then aligned to the assembled contigs using Bowtie 2 v. 2.3.3 [103]. Inconsistencies were identified and polished using Pilon [104]. Contigs smaller than 500 bp and with less than 2.0 k-mer coverage were filtered out. Quality of genomes were assessed with CheckM [105]. Assembled genomes were annotated using the IMG/JGI platform [106]. The genome data generated for this study are available in NCBI BioProject PRJNA941448.

### Core gene phylogeny

In a previous study, we defined a set of 1,356 ‘core genes’ from 58 genomes of *Xep* strains isolated from Florida [47]. The core genes were determined based on amino acid sequence homology using GET_HOMOLOGUES software package [107]. We used the core genes from a representative *Xep* genome, Xp91-118, as query to search the remaining 269 genomes using local BLAST [108]. BLAST results were filtered using query coverage and pairwise nucleotide sequence alignment thresholds of 70% each and the sequence was checked for the presence of standard start and stop codons at either end of the gene and gene was removed if both were not present. A total of 887 genes were found to be intact in all 270 genomes. Genes were individually parsed and aligned using MAFFT [109] and concatenated using sequence matrix [110]. The result was a 617.854 Kbp alignment, hereafter referred to as core genes.

The concatenated core gene sequence was used to construct a maximum likelihood (ML) phylogenetic tree using RAxML v.8.2.12 [111]. General time reversible model with gamma distributed rates and invariant sites (GTRGAMMA) was used as the nucleotide substitution model. To account for recombination, the ML tree output from RAxML and concatenated core genome alignment were used as was input for ClonalFrameML v1.12 [70].

### Population structure

SNPs were extracted from core genes for hierarchical clustering based on Bayesian analysis of population structure (hierBAPS) algorithm [112], implemented in the ‘rhierBAPS’ R package v 1.0.1 [71, 113]. For visualization, hierBAPS clusters were added to the phylogenetic tree generated from ClonalFrameML using the ‘ggtree’ package in R [114]. The treemap function in plotly [115] was used to show the relative distribution of clusters across geographic locations. R package ‘ggplot2’ was used to map hierBAPS clusters to countries [116]. The ‘PopGenome’ R package [117] was used to calculate FST, nucleotide diversity (pi), and Tajima’s D statistic by geographic location and by hierBAPS cluster.

Assembled genomes were used for calculating average nucleotide identity and pangenome analysis. Average nucleotide identity (ANIb) between strains was calculated using assembled genomes with Pyani version 0.2.10 [118]. The pangenome was estimated using Roary v3.12.0 [119] after annotation from Prokka v1.12 [120]. The gene presence absence matrix from Roary (S4 Data) was used as input for generation of NMDS plots using the ‘dplyr’ and ‘ggplot2’ packages from tidyverse [116] and to generate gene accumulation curves for each cluster using package ‘micropan’ [121].

### Bayesian analysis of *X. euvesicatoria* pv. *perforans* divergence times

A whole genome alignment was generated using split k-mer analysis version 2 (SKA2) [122] for all 270 *Xep* strains plus outgroup *X. euvesicatoria* pv. *euvesicatoria* strain 85-10 (NCBI Accession GCA_000009165.1; S2 Data). The alignment was reduced to variable sites only using Geneious 2023.2.1 (BioMatters Ltd.) A phylogenetic network was calculated from the resulting SNPs using the NeighborNet 2004 algorithm in SplitsTree5 [123, 124]. Phylogenetic conflict was indicated between the 259 strains, cluster 10 strains, and outgroup (S1 Figure, part B). Removing the cluster 10 strains did not remove the conflict between *Xep* and *Xee* outgroup. As a result, we limited our dating of the phylogeny of *Xep* to the 259 strains in BAPS clusters 1 through 9. We used Gubbins v. 2.4.1 [72] to remove putative recombinant sites from whole genome alignments generated using SKA2 [122] and the complete genome of Xp91-118 as a reference (GCF_000192045.2). The resulting alignment was used to infer a phylogenetic tree using the GTRGAMMI model in RAxML version 8.2.10 [125]. The temporal analysis was conducted with BactDating v1.1.1 [73]. The inputs to the BactDating analysis were the maximum likelihood tree and dates of isolation assigned as dates of tips. The rooting of the tree was estimated using the initRoot function, which maximizes the correlation between tip date, the year the strain was collected, and root-to-tip branch lengths. Dates of nodes were inferred using the bactdate function on the re-rooted tree using a relaxed molecular clock with Markov chain Monte Carlo (MCMC) chains of 10^6^ iterations. Phylostems [126] was used to assess the temporal signals within internal clades for interpretation of node date inferences.

We also used BEAST v. 1.10.4 (Suchard et al. 2018) to infer a dated phylogeny. The XML file was manually edited to include the ‘ascertained’ flag in the alignment block (S3 Data). The HKY nucleotide substitution model with empirical base frequencies and gamma distribution of site-specific rate heterogeneity was used with coalescent Bayesian skyline priors with an uncorrelated relaxed clock for Bayesian phylogenetic inference over MCMC chains of 200 million generations. Adequate mixing was assessed based on a minimum effective sample size of 200 for parameter estimates as calculated by Tracer v. 1.10.4. A maximum clade credibility tree was inferred from the posterior distribution of trees using TreeAnnotator v. 1.10.4, specifying a burn-in of 10% and the ‘keep’ option for node heights. Trees were visualized in iTOL version 6.9.1 [127].

### Type III Effector Analysis

A T3E effector database was curated using amino acid sequences of 66 *Xanthomonas* effectors [128] (S6 Table). When available, functional annotations were retrieved from NCBI and Pfam databases [129]. Orthologous sequences were identified with the software BLASTp [130, 131], by querying the curated effectors database against the amino acid sequences of the annotated genomes of 270 *Xep* strains. Sequences (BLAST hits) were considered effector orthologs when at or above a threshold of 70 percent identity and 50 percent query coverage. When multiple sequences from the same strain had hits above the thresholds to a particular effector, a weighted calculation of the identity and coverage was used to select the best hit. Sequences with homology to multiple effectors and sequences with evidence of contig breaks were manually removed. Assignment of sequences as effector orthologues was confirmed by performing a clustering analysis of all sequences using the software USEARCH v. 11.0.667 and the algorithm HPC-CLUST [132]. For the duplicated effector XopP, we used a phylogenetic analysis of all sequences to distinguish likely orthologous alleles from the more genetically distant paralogous sequences, which were assigned to XopP2.

Orthologous sequences from each effector were extracted from the annotated genomes, aligned with MAFFT [109], and allelic variants identified [133] to generate a numeric matrix representing presence and allelic variant or absence. Hierarchical clustering analysis of effectors was performed by calculating a distance matrix with function ‘dist’ with the method ‘manhattan’, and the function ‘hclust’ with the method ‘complete’ from the R package ‘vegan’ [113, 134]. The results were displayed as a heatmap with the package ‘gplots’ and the function heatmap.2 [135].

To investigate the presence of positive selection acting on the effector sequences, we used the software HyPhy (Hypothesis Testing using Phylogenies) implementing the methods FUBAR (Fast, Unconstrained Bayesian AprRoximation) and MEME (Mixed Effects Model of Evolution) [77, 78]. The Bayesian method FUBAR evaluates pervasive selection, assuming the same rates of synonymous and nonsynonymous substitution per site on all branches. The method MEME uses a maximum likelihood approach to evaluate episodic selection, i.e., selection only a subset of branches of the phylogeny. For each effector gene, a codon-aware alignment was generated with the software PRANK using the codon flag ‘-c’ as settings [136]. RAxML [111] was used to infer a phylogenetic tree with the GTRGAMMA (gamma time-reversible) model of nucleotide substitution. The codon-aware alignment and phylogenetic tree were used as the input files for FUBAR and MEME.

To determine the relationship of the effector profiles with respect to core gene cluster, geographic and temporal distribution, we transformed the dissimilarities in the matrix of effector profiles into distances with non-metric multidimensional scaling (NMDS). We used the Bray-Curtis dissimilarity index, a robust index able to handle missing data that considers the presence and absence of effectors as equally informative, calculated with the package ‘vegan’ and the function ‘metaMDS’ [134]. We used a low number of dimensions (K=2) and set try=30 and trymax=500 for random starts to avoid the NMDS getting trapped in local optima. NMDS plots were created with the packages ‘ggrepel’ and ‘ggplot2’ [137, 138]. Based on the NMDS analysis, we assigned strains to effector clusters, which were plotted on a worldwide map with the packages ‘ggplot2’ and ‘scatterpie’ [138, 139]. The map was created in R with the packages ‘cowplot’, ‘ggrepel’, ‘ggspatial’, ‘libwgeom’, ‘sf’, ‘rgeos’, ‘memisc’, ‘oz’, ‘maptools’ and ‘rnaturalearth’ with the function ‘ne_countries’ [137, 140–147]. Geographic coordinates (longitude, latitude) of countries and states (for USA) of collection were obtained with the R package ‘googleway’ [148] and the function ‘mutate_geocode’ from Google maps.

To sequence the putative TAL effector from 2P6S1, native plasmid DNA was isolated using the alkaline lysis method [149]. *Eco*RI digested DNA of the plasmid prep was ligated into vector pLAFR3 [150] restricted with the same enzyme for transformation into *E*. *coli* DH5α. Clones containing the TAL effector were identified by PCR and analyzed by restriction digest. One clone, designated as p7.1, contained an approx. 5 Kbp *Eco*RI fragment and was selected for Sanger sequencing and phenotype testing. For Sanger sequencing of the TAL repeat region, DNA of p7.1 was restricted with *Nsi*I and the internal fragment was ligated into vector pBluescript restricted with *Pst*I. Additional pBluescript subclones were made using *Bam*HI (∼3 Kbp and ∼1.1 Kbp) and *Bam*HI/*Eco*RI (∼1 Kbp) in order to cover the entire cloned region in p7.1. All clones were transformed into DH5α for sequencing using vector primers T3 and T7.

Copper resistance genes in assembled genomes were identified with BLASTn analysis using *copL* (MBZ2440241.1), *copA* (MBZ2440240.1), and *copB* (MBZ2440239.1) from *Xep* strain Xp2010 as reference sequences [30].

## Acknowledgements

We thank D. Ritchie and C. Mauney for providing reference strains and collecting disease samples, respectively. Tina Simonton and Phyllis May provided assistance with strain collection in Ontario. This research was supported by USDA NIFA awards 2015-51181-24312, 2020-67013-31921, and 2022-51181-38242, Ontario Agri-Food Innovation Alliance, Ontario Tomato Research Institute, and the Fresh Vegetable Growers of Ontario, the North Carolina Tomato Growers Association. Plant diagnostic laboratories receive funding from USDA-NIFA through the National Plant Diagnostic Network. FIB was supported by Conacyt Mexico. MK was supported by Swedish International Development Cooperation Agency.

## Author Contributions

ST, FIB, MOJ, PDR, GEV, JBJ, and EMG conceived the study. MOJ, GVM, PA, TBA, GC, LBTdlB, EB, TCC, DTKT, TAC, DSE, RFG, DMF, MK, MLI, FJL, LL, ETM, SAM, NTTN, EO, AMQD, RR, FR, GER, VMS, PT, CT, JBJ, and GEV collected bacterial strains. ST, MOJ, GVM, and JKG prepared genomic libraries. ST, FIB, MOJ, JHT, A Sharma, A Subedi, AK, and EMG performed data analysis. ST, FIB, MOJ, A Sharma, A Subedi, GEV, JBJ, and EMG drafted the manuscript. All authors reviewed the manuscript.

## Data summary

The raw read files and genome assemblies for all strains sequenced in this study are deposited in NCBI under BioProject PRJNA941448. The sources of genome assemblies acquired from public databases are listed in Table 1.

## Supporting Information

**S1 Figure.**
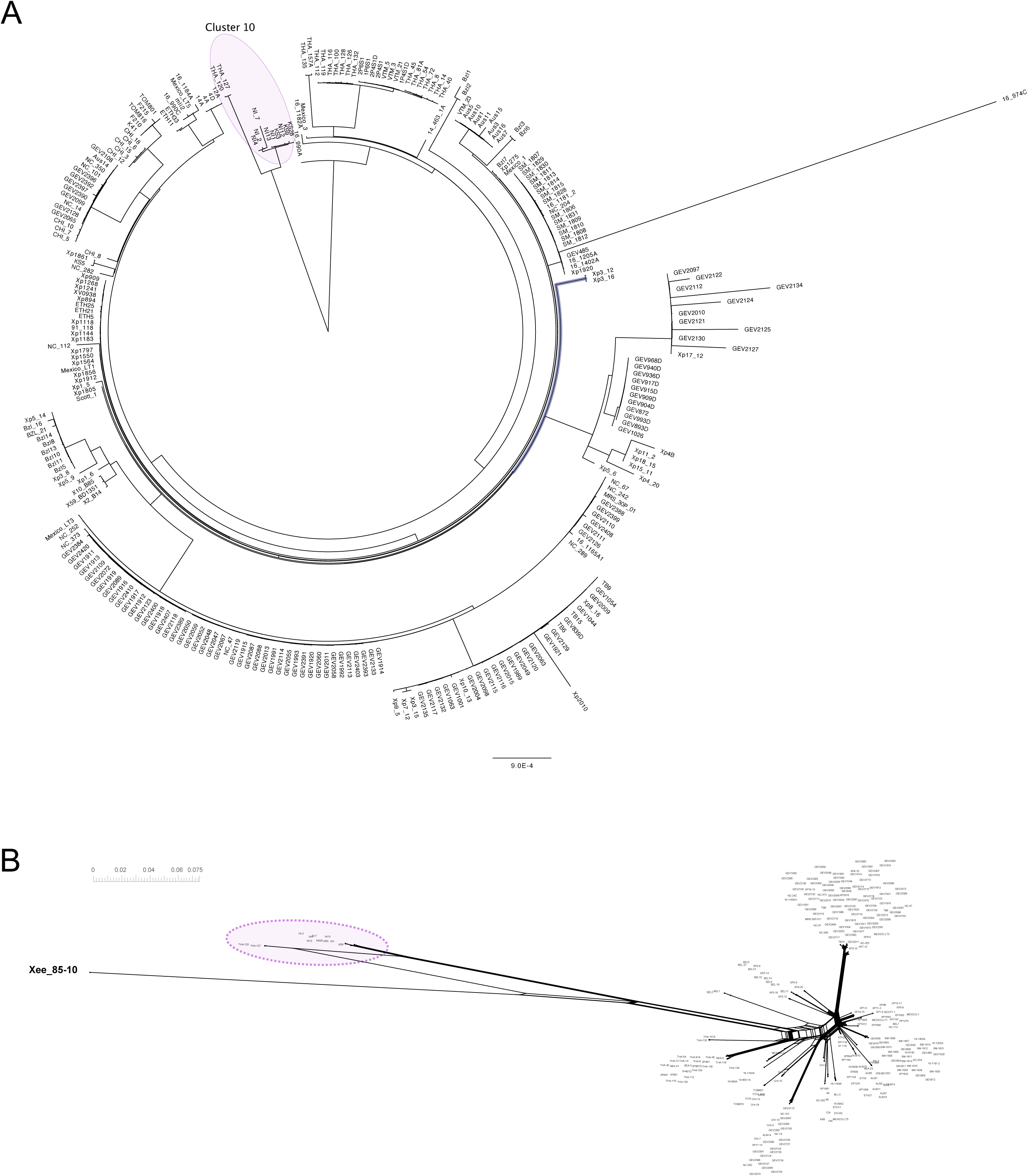
Phylogenetic analysis of 270 *X. euvesicatoria* pv. *perforans* strains. (A) Maximum likelihood phylogenetic tree of *Xanthomonas euvesicatoria* pv. *perforans* strains based on aligned nucleotide sequences of 887 core genes. The tree was inferred using RAxML using a GTRGAMMAI substitution model. The tree was rooted using the 11 genetically diverged strains that make up core gene cluster 10. (B) Neighbor-net network inferred using SNPs from aligned whole genome sequences, including *X. euvesicatoria* pv. *euvesicatoria* strain 85-10 (bolded) as an outgroup. Core gene cluster 10 strains are highlighted. Reticulations in the network indicate conflicting phylogenetic relationships.

**S2 Figure.**
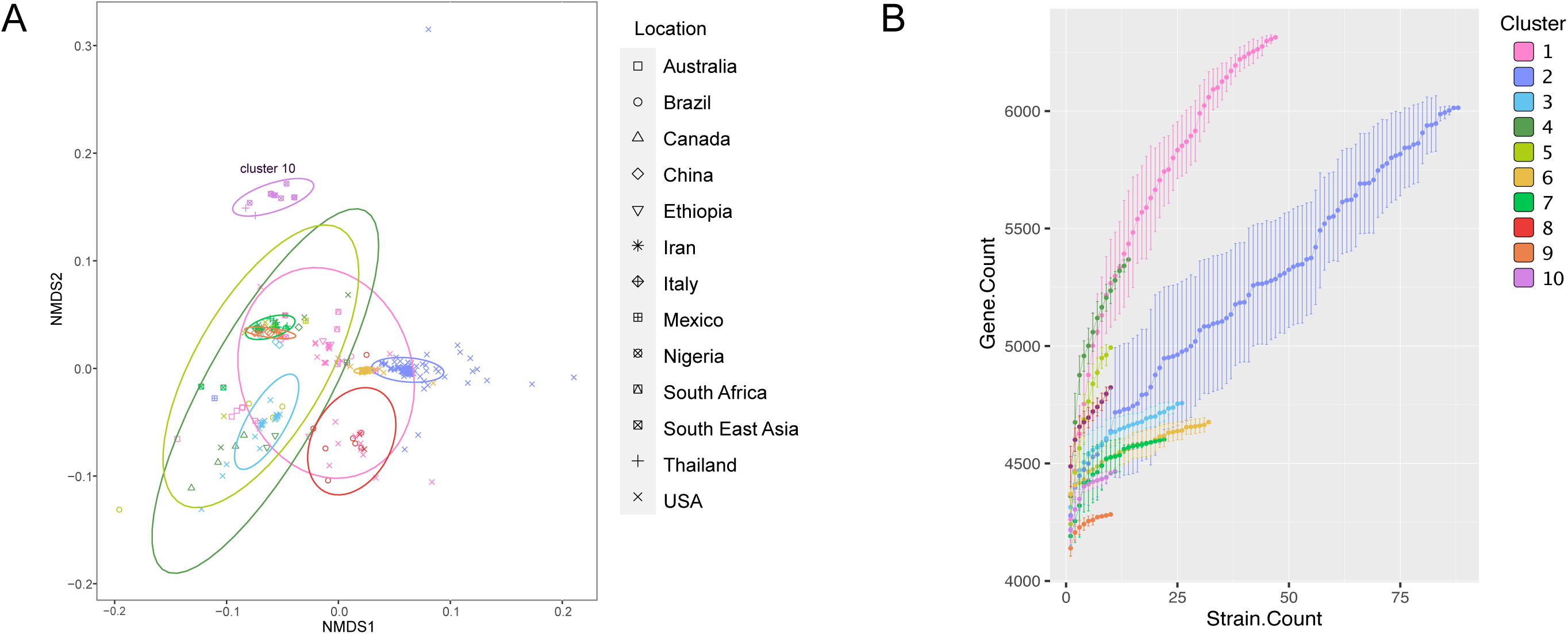
Accessory genome variation in *Xanthomonas euvesicatoria* pv. *perforans*. (A) Visualization of pangenome variation by non-metric multidimensional scaling of gene presence-absence for all 270 *X. perforans* strains by BAPS cluster. Ellipses assume a multivariate t-distribution. (B) Increase in gene count with increasing number of strains sampled. Clusters 1 and 2 were represented by the most strains, but other clusters showed similar rates of increase in the pangenome of the cluster.

**S3 Figure.**
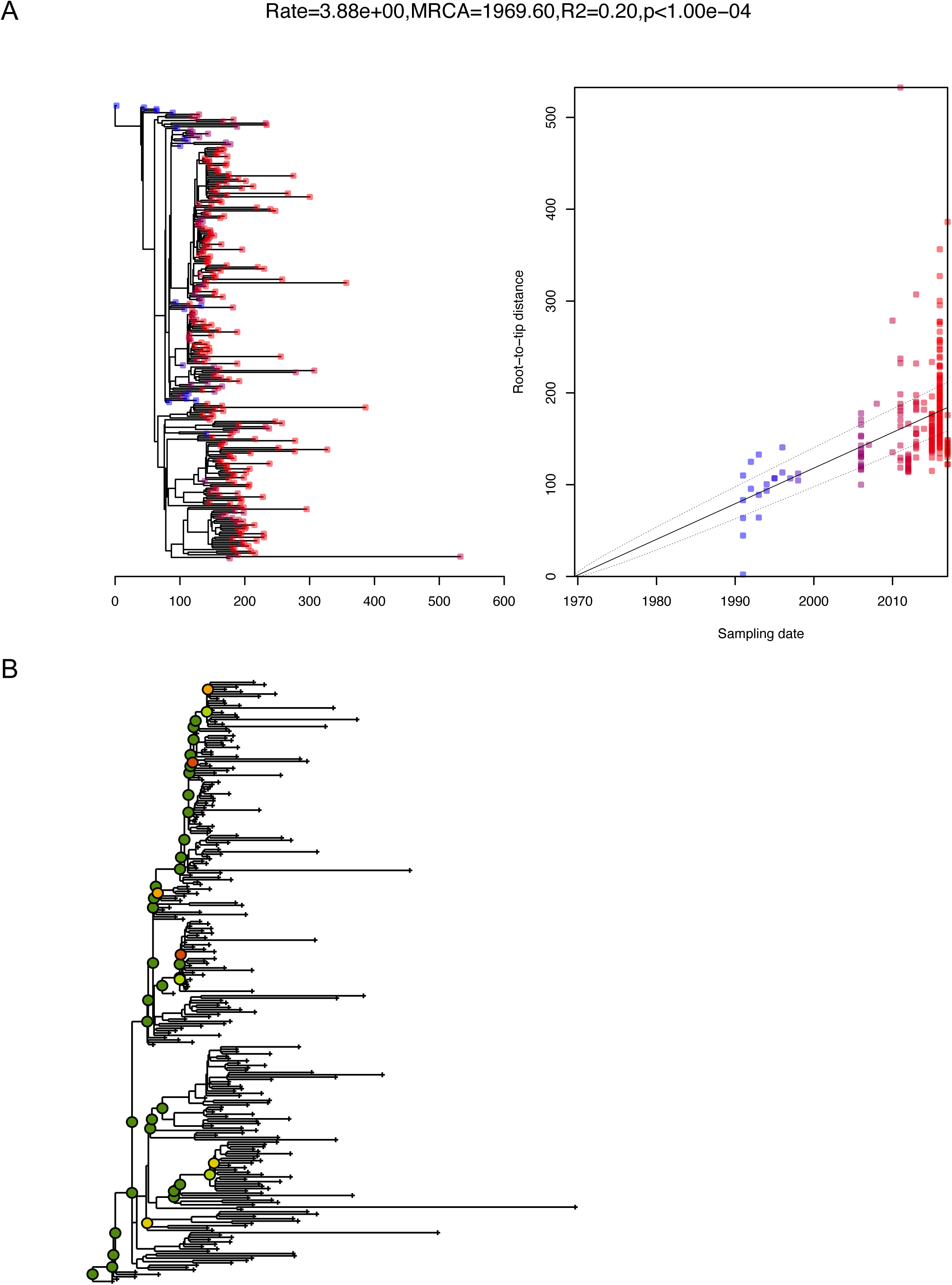
Temporal signal in phylogenetic tree of 259 *X. euvesicatoria* pv. *perforans* strains. (A) Correlation between sampling year and root-to-tip distance in maximum likelihood phylogenetic tree inferred from alignment of whole genome sequences. Output was generated from BactDating R package. (B) Temporal signal within the phylogenetic determined using Phylostems tool. Tree is rooted as in Figure 2. Nodes with statistically significant temporal signals are indicated with colored circles. Adjusted R-squared values by color: dark green 0–0.2; light green 0.2–0.4; yellow 0.4–0.6; orange 0.6–0.8; red 0.8–1.

**S4 Figure.**
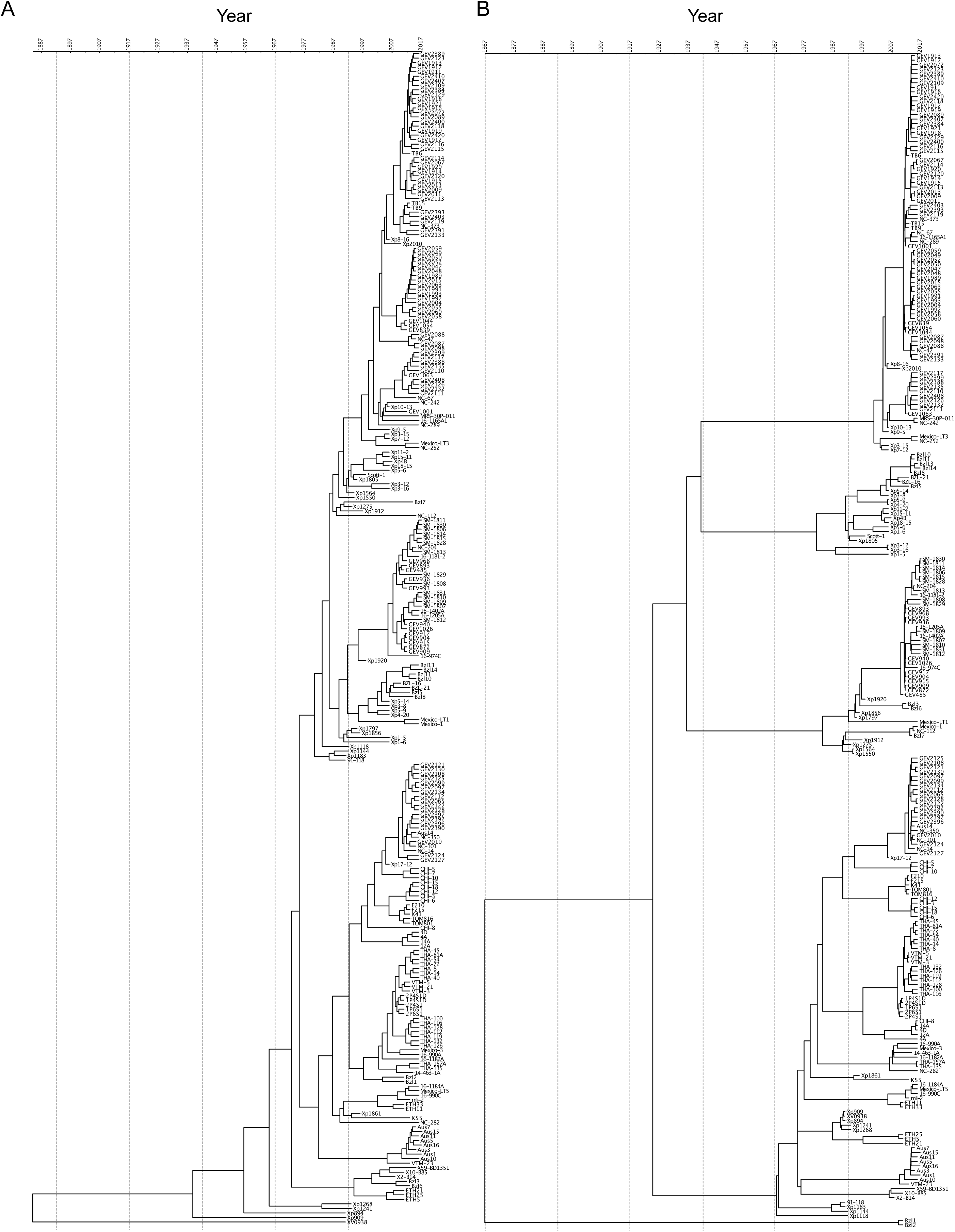
Dated phylogenies of 259 *X. euvesicatoria* pv. *perforans* strains. (A) Dating using BactDating relaxed clock analysis on RAxML-generated phylogeny. This is the tree shown in Figure 2, shown here without collapsed nodes. (B) Dating of same dataset using BEAST with coalescent Bayesian skyline priors and an uncorrelated relaxed clock.

**S5 Figure.**
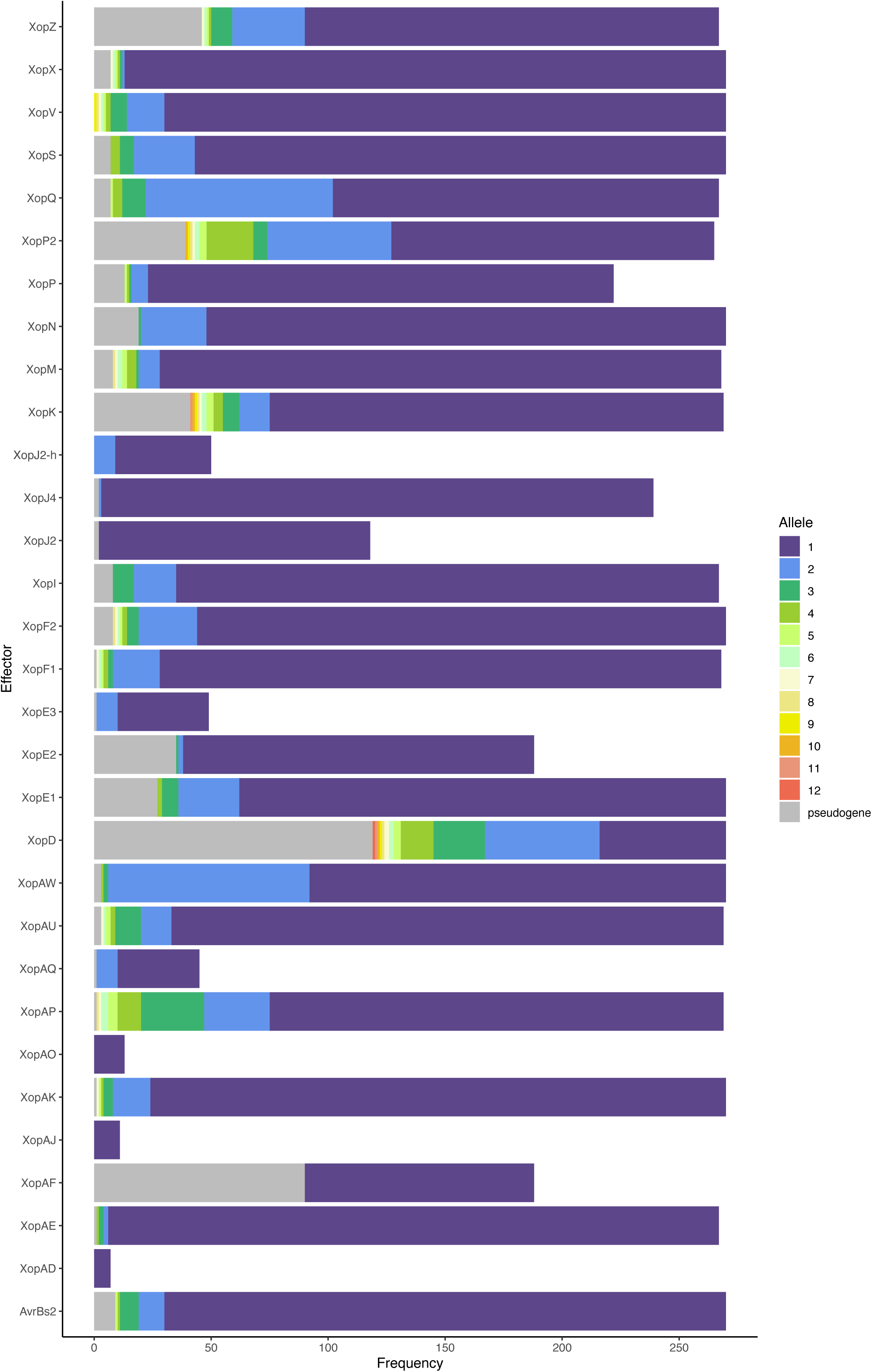
Frequency of Xop effectors among 270 *Xanthomonas euvesicatoria* pv. *perforans* strains. The most common allele observed was assigned to allele type 1, second most frequent allele to allele type 2, and so on. Note that alleles classified as pseudogenes included contig breaks, which include assembly errors. For example, all strains appear to have *xopD*, but a repeat caused a contig break in the gene in nearly half of the genomes.

**S6 Figure.**
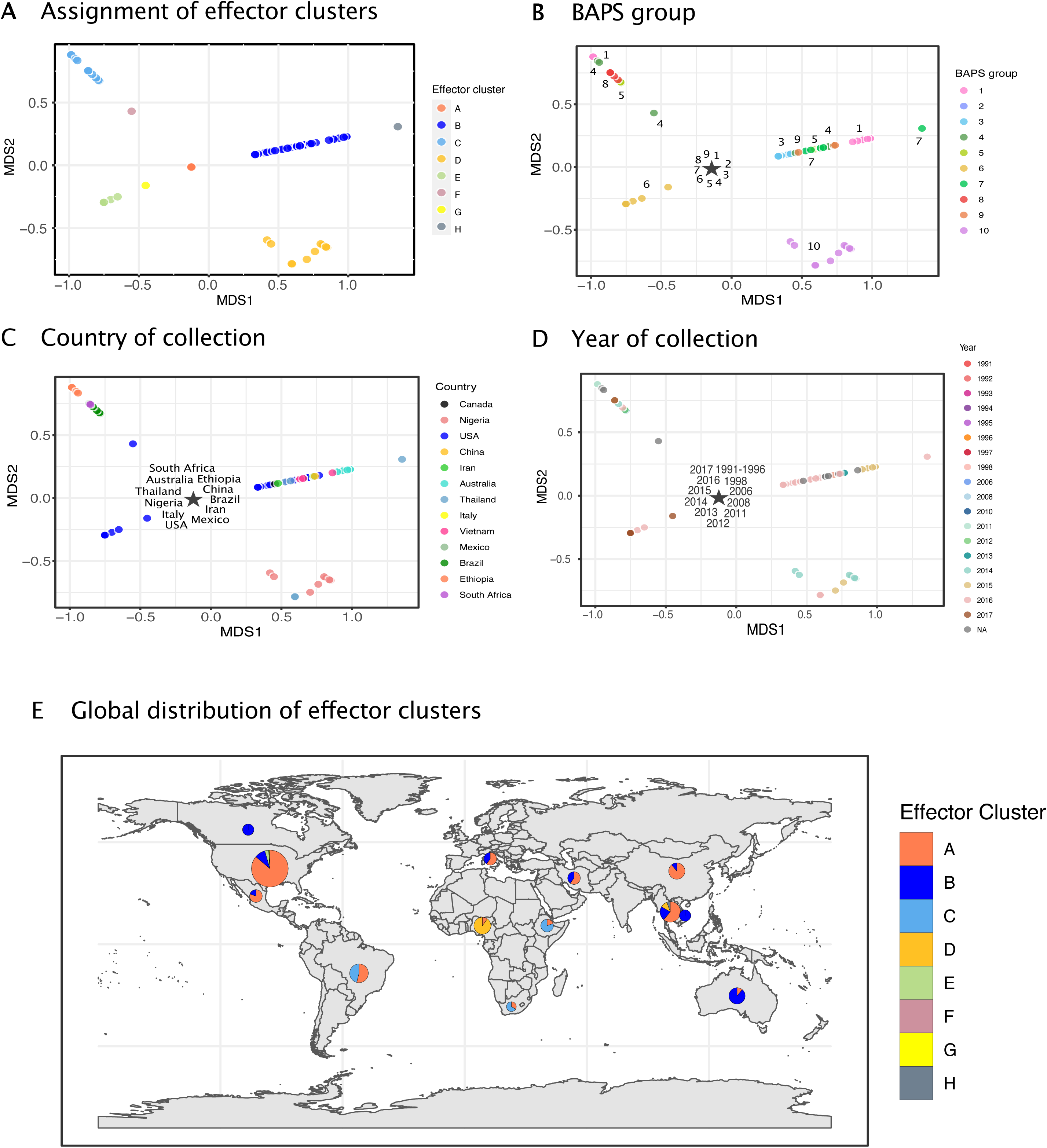
Clustering of 270 *Xanthomonas perforans* effector profiles by non-metric multidimensional scaling and distribution of resulting clusters among geographic regions. Analysis did not include TAL effectors. (A) The most frequently observed group of effector profiles form cluster A. This cluster of 188 strains is represented as a star in plots B-C, as it is represented in most BAPS core gene clusters (B), most of the sampled tomato production regions (C), and in collections from 1991 to 2017 (D). Clusters were largely defined by low frequency effectors (S7 Figure). (E) Distribution of strains by effector clusters among sampled countries.

**S7 Figure.**
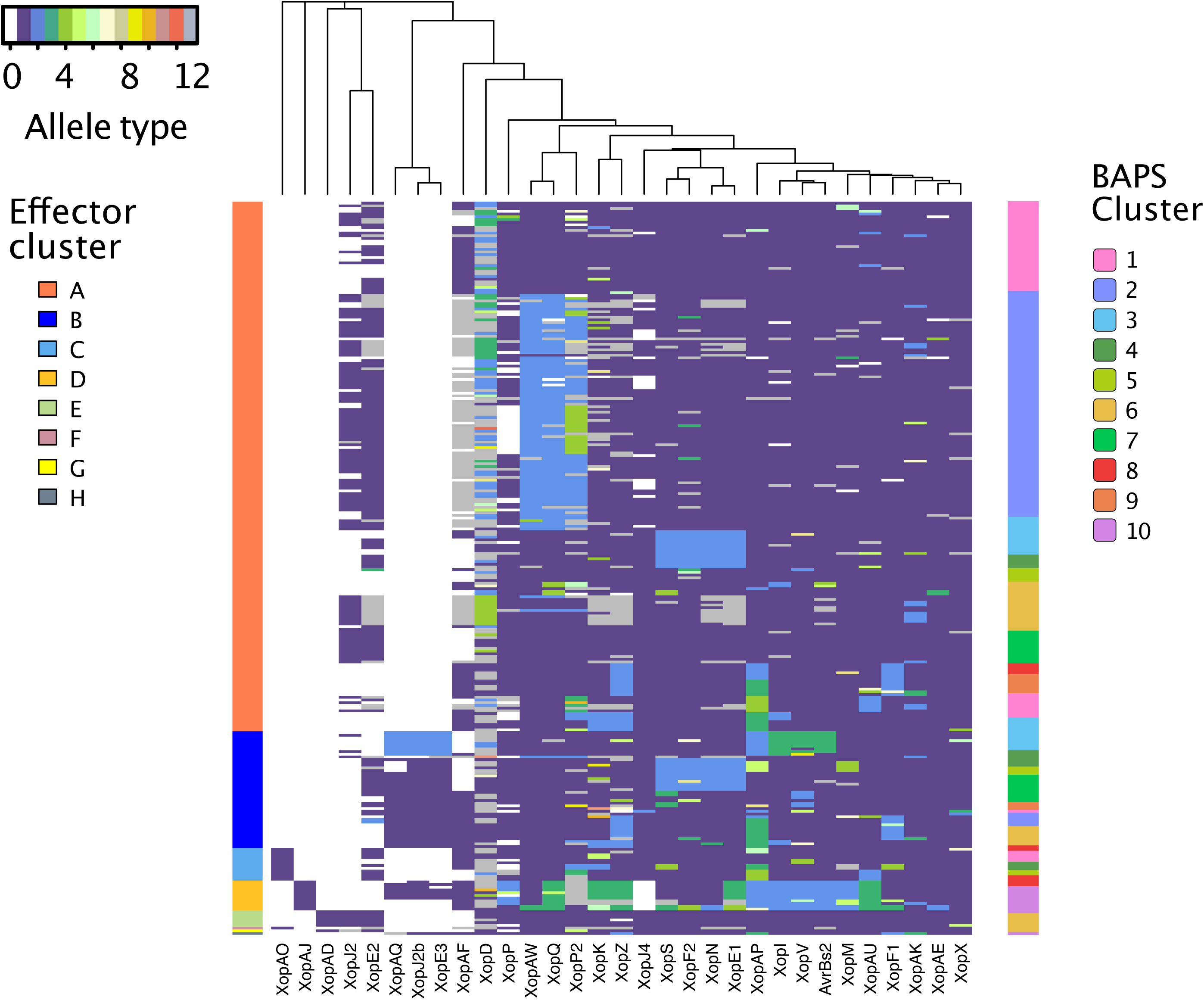
Variation in Type III secreted effector profiles in 270 *Xanthomonas euvesicatoria* pv. *perforans* strains ordered according to NMDS of effector profiles. Analysis did not include TAL effectors. Type III secreted effectors are in columns and *X<u>ep</u>* strains in rows. Effector status is shown by allele type: absence is indicated by allele type 0 (white), while the most frequent allele observed when the effector is present is allele type 1 (purple), second most frequent is allele type 2 (blue), and so on. Putative pseudogenized effectors are shown as allele 13 (gray). The order of columns was determined by hierarchical clustering analysis, placing similarly distributed effectors adjacent to each other. Order of rows is based on NMDS clustering analysis of effector profiles (see S6 Figure, part A).

**S1 Table. Genome data and metadata for *X. euvesicatoria* pv. *perforans* strains.** (A) BAPS cluster assignment for each strain and NCBI information for each genome. Genome assembly statistics are given for newly sequenced strains. (B) Depth of coverage relevant to ANI comparisons in Fig. 1D.

**S2 Table. Genetic diversity statistics by geographic region and BAPS group.** (A) Statistics by country and U.S. state. (B) Statistics by BAPS group.

**S3 Table. Average nucleotide identity (ANIb) comparisons between strains with highly similar core gene sequences collected across continents.** (A) Proportion nucleotide identity. (B) Alignment fraction.

**S4 Table. Putative type III effectors (Xop proteins) found in 270 *X. euvesicatoria* pv. *perforans* assembled genomes.** (A) Summary for each locus. (B) Results by strain. Each different amino acid sequence per gene was assigned a numerical allele type, such that the most common allele observed was assigned to allele type 1. Potential pseudogenes are indicated with “pseudo” and absence indicated with zero. Locus tags refer to JGI IMG annotations (https://img.jgi.doe.gov). Reference sequences used for BLAST searches are given in S5 Table. The final column shows the result of BLAST searches for TAL effectors.

**S5 Table. Presence or absence of copper genes (*copLAB*) in 270 *X. euvesicatoria* pv. *perforans* assembled genomes.** Symbols represent gene presence ‘+’ or absence ‘-’. Contig break in gene is indicated by (+).

**S6 Table. Type III effector database used to query assembled genomes for effector genes.**

**S1 Data. Nucleotide alignment of 887 core genes from 270 *X. euvesicatoria* pv. *perforans* strains.** Alignment is 617,855 bp in FASTA format.

**S2 Data. Nucleotide alignment of variable sites from whole genome alignment of 270 *X. euvesicatoria* pv. *perforans* strains and *X. euvesicatoria* pv. *euvesicatoria* strains 85-10.**

**S3 Data. XML file used for BEAST analysis.**

**S4 Data. Pangenome matrix for 270 *X. euvesicatoria* pv. *perforans* strains**

